# A preliminary study of the cytotoxicity of the protein extract of *Chrysobalanus icaco* L. and *Eugenia astringens* Cambess., commercialized in markets

**DOI:** 10.1101/720052

**Authors:** Thaís A. Pedrete, Julianderson O.S. Carmo, Emiliano O. Barreto, Josino C. Moreira

**Author notes:** Department of Physiological Sciences, Institute of Biology Roberto Alcantara Gomes – State University of Rio de Janeiro (UERJ), Rio de Janeiro, RJ, 20550-170, Brazil. Corresponding author (Thaís A. Pedrete).

## Abstract

The use of plants and their products for medical treatment is very common procedure in Brazil, especially for treatment of diabetes. In fact, several plants can demonstrate hypoglycemic effects in vitro assays. However, the use for human treatment requires the knowledge of their toxicological properties. The aim of this study was to evaluate the effect of protein extracts of *Chrysobalanus icaco* collected from natural habitats and of *Eugenia astringens* acquired from the market of Rio de Janeiro on the viability and migration of fibroblasts. *E. astringens* has a similar morphology as *C. icaco* and it is sold as *Chrysobalanus* in a popular market in Rio de Janeiro. Being a different plant, *E. astringens* expresses different proteins, and its protein extract has proved to possess higher toxic properties than *C. icaco* does. Cytotoxicity assays indicated that, as the protein extract concentration increases, fibroblast viability decreases. Only the *E. astringens* extract displayed cytotoxicity at all concentrations, in addition to reduced fibroblast migration. The results obtained in this study demonstrates that it’s necessary integrative policies for rational use of medicinal plants and their commercialization, since the current use of medicinal plants may be inadequate and it is of great importance for Public Health.

## 1. Introduction

Several plants are widely used for medical purposes by the population, but this use is most often made from a lay indication, without knowing the risks of toxic effects. Besides, there is no guarantee of the provenance and proper storage of these supposedly “medicinal plants”. It is clear that there is a lack of incentive and scientifically-based information to integrative and complementary practices and actions to promote the safe and rational use of medicinal plants, including information on how the species should be prepared and used by the population (Bochner *et al*., 2012).

The leaf extract (tea) of the plant *Chrysobalanus icaco* L., a species of restinga popularly known as abajerú, is used in folk medicine because it exerts biological activities, such as the decrease of blood sugar levels, being indicated for the treatment of diabetes, besides be diuretic and antioxidant (Venancio *et al*., 2018). Another plant (*Eugenia astringens*, Cambess.) which is morphologically similar to *C. icaco*, also is known by the same popular name of abajerú and is commercialized as *C icaco* (Bochner *et al*., 2012; Silva and Peixoto, 2009). These two species may not possess the same therapeutic and toxicological properties, which are of concern to Public Health. The attribution of hypoglycemic activity to *E. astringens* may indicate a misconception since other species of Myrtaceae have hypoglycemic potential (Silva and Peixoto, 2009). So, in order to clarify the toxicological aspects of the extract obtained from these 2 plants, a cytotoxic assay was performed.

For cytotoxicity studies in animal cells several techniques, using distinct cell types as a target, are available. Cytotoxicity means the determination of any toxic effecs at the cellular level, such as changes in membrane permeability, cell death or enzymatic inhibition resulted from exposure to a toxicant, in this case, the studied plants or plant products (Stockert *et al*., 2012).

Cell viability can be evaluated by several methods, among which the one which involves the conversion of 3-(4,5-dimethylthiazol-2-yl)-2,5-diphenyltetrazolium bromide (MTT) to formazan by mitochondrial reactivation in active-living cells (Zandi *et al*., 2016). The MTT assay is a standard colorimetric assay that estimates the cytotoxic potential of the samples, in addition to measuring the cellular proliferation of drug agents. Cell viability is expressed as a percentage of live cells from the tested material, comparing with the percentage of cells of the cytotoxicity positive control (Stockert *et al*., 2018).

Another test to evaluate the toxicity of the plant extract is the Scratch Wound Healing Assay, which allows measuring the migration of cells that is a phenomenon present in the healing process. It is a method in which a crack imitates a wound in a monolayer of confluent cells so that the cells at the edge gradually move towards the crack (Manoj *et al*., 2009). Wound healing is the process of repairing and regenerating the dermis and epidermis that accompanies the lesions (Liang *et al*., 2007; Pitz *et al*., 2016). The evaluation of the healing activity of plant extracts is scarce at the cellular level. Fibroblast cultures have been proposed as a method for the investigation of wound healing activity, since these cells are the main source of extracellular connective tissue matrix and the migration of fibroblasts is considered vital for rapid and effective skin repair damaged (Manoj *et al*., 2009).

The lack of data about the toxicity of the protein extract of these 2 plants commercialized as abajerú (*C. icaco* and *E. astringens*), protein extracts of *Chrysobalanus icaco* collected directly from its natural habitats and of *Eugenia astringens* acquired from the market of Rio de Janeiro was performed using the viability and migration of fibroblasts assay.

## 2. Material and methods

### 2.1 Plant sampling

*Chrysobalanus icaco* leaves were collected directly from its natural habitats, Praia Grande – Arraial do Cabo-RJ (PG; -22,9696606, -42,0302859), Restinga de Massambaba – RJ (RMA; -22,9337727, -42,4267012), Marechal Deodoro – AL (AL; - 9,7823233, -35,852364), as shown in the map (Fig. 1). *Eugenia astringens* leaves were purchased on the market Mercadão de Madureira located in the North zone of Rio de Janeiro city.

**Fig. 1.**
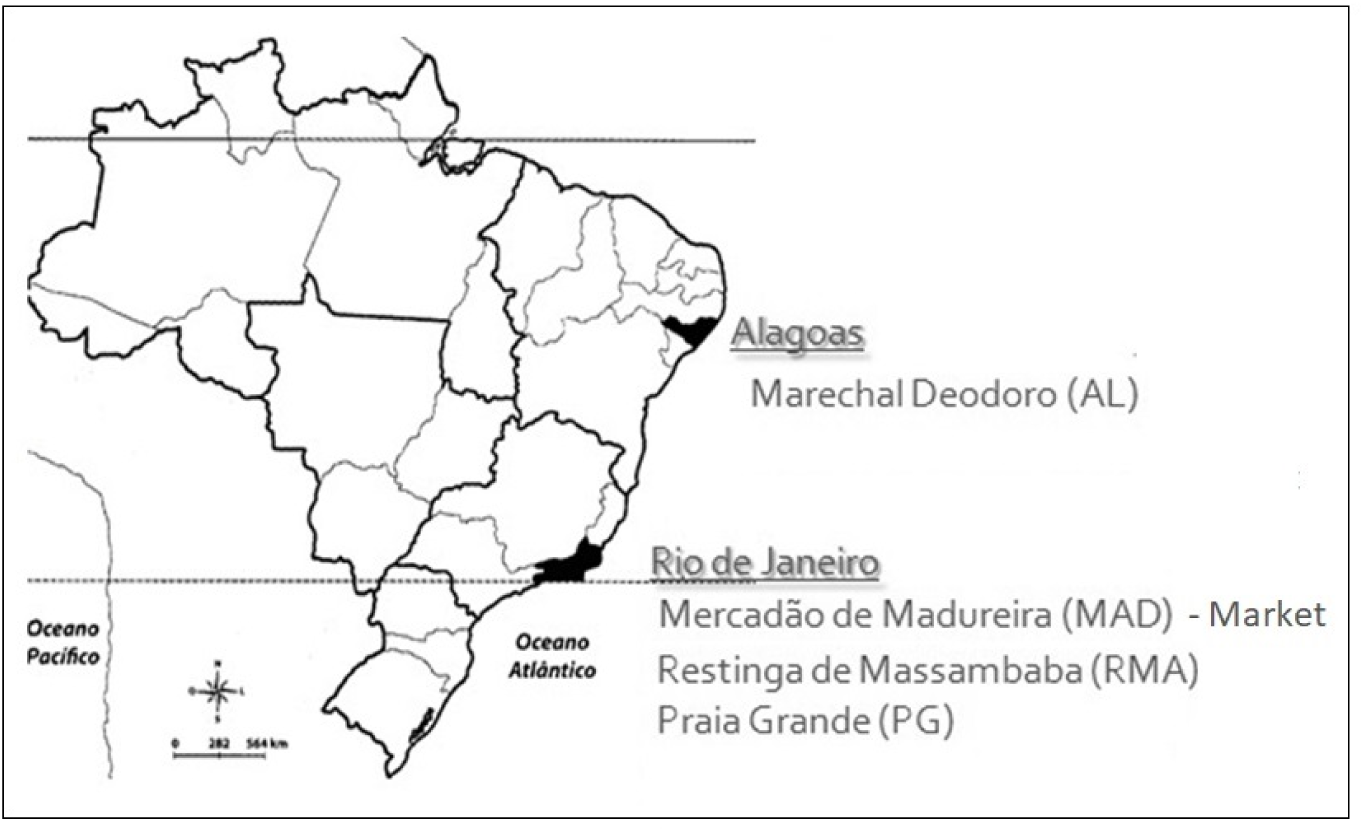
Sampling sites. AL - Marechal Deodoro, MAD – Mercadão de Madureira, RMA - Restinga de Massambaba, PG - Praia Grande (PG).

### 2.1 Protein extraction

About 10 mg of each lyophilized sample were weighed into microtubes in triplicate. Samples were incubated in the presence of 400 μl lysis buffer (4% SDS 0.1 M Tris-HCl buffer pH 7.6) at 95 °C for 15 min in a thermomixer. The lysate extract was frozen at -80 °C for further quantification of total proteins by the Lowry method (Lowry et al., 1951), using bovine serum albumin (2.0 mg/mL) as the standard for the analytical curve. Samples (2 μL) and analytical curve (0, 10, 20, 30, 40, 50, 60 and 70 μg/mL) were read in a Jasco V-530 spectrophotometer at the wavelength of 750 nm.

### 2.2 Cytotoxicity evaluation of protein extracts

This assay was performed as follows:

#### 2.2.1 Cell culture

Fibroblasts (3T3 cell line) were kept in medium Dulbecco’s Modified Eagle Medium (DMEM), containing 10 % fetal bovine serum, L-glutamine (2 mM) and gentamicin (40 μg/mL) in incubator at 37 °C and 5 % CO_2_.

##### 2.2.1.1 Cell viability assay

The effect of the protein extracts of *Eugenia astringens* (Mercadão de Madureira) and *Chrysobalanus icaco* (Restinga de Massambaba - RJ, Marechal - AL, Praia Grande - RJ) on fibroblasts viability was evaluated through the MTT assay (Mosmann, 1983).

The cells were seeded in a 96 well plates and placed in CO_2_ incubator overnight. The cells were then treated with different sample solutions (1, 5, 10 and 20 μg/mL) in four replicates. The control group was treated only with the medium (DMEM). After treatment, a solution of MTT (3-(4,5-dimethylthiazol-2-yl)-2,5-diphenyltetrazolium bromide) (5 mg/mL in phosphate buffered saline - 1X PBS pH 7.4) was added to each well and incubated for 4 hours. Subsequently, the supernatant was discarded and 150 μl of dimethyl sulfoxide were added for solubilization of the formazan crystals. The absorbance was measured using a microplate spectrophotometer (DTX 880 Multimode Detector, Beckman Coulter), adjusted to 595 nm, and the optical density was calculated (Equation 1).

Equation 1 – Optical density of cells submitted to the cell viability assay.

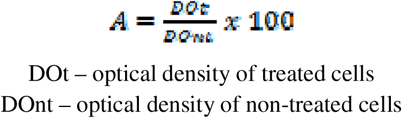

##### 2.2.1.2 Scratch wound healing assay

The effect of the protein extracts of *Eugenia astringens* (Mercadão de Madureira), *Chrysobalanus icaco* (Restinga de Massambaba - RJ, Marechal - AL, Praia Grande - RJ - branch) on fibroblast migration was evaluated through cell migration technique, method described by Liang *et al*. (2007).

Cells (7 × 10^4^ cells / well, measured by the Newbauer’s chamber) were seeded in 24-well plates and maintained overnight for cell adhesion and formation of a monolayer at approximately 80% confluency. A small part of the monolayer was removed in the middle of the plate with a 200 μL pipette tip (a scratch is placed on the monolayer and the part removed is discarded). Cells were washed with phosphate buffered saline and treated with 5 μg/mL of the samples or culture medium (control) in triplicate. This exposure concentration at which some effects started to be observed in the cell viability assay was chosen to perform the present assay. Cell migration was assessed by photomicrographs at 0- and 24-hours post-exposure using an inverted microscope (Olympus IX70) with digital camera to measure the area of wound closure. The photomicrographs were analyzed using Image J software and cell migration was expressed as the area in pixels, so that the percentage of closure of the initial area formed was determined quantitatively (Equation 2).

Equation 2 – Migration rate of fibroblasts submitted to the cell migration assay.

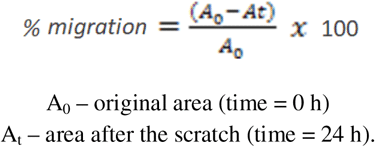

### 2.4 Statistical analysis

The results of the cell viability and migration tests were expressed as mean ± standard error, performed in triplicate and analyzed statistically using analysis of variance (ANOVA), followed by Newman-Keuls post-test. The results were considered significant when p <0.05. All results were analyzed using GraphPad Prism® software version 5.01 (GraphPad Software Inc, San Diego CA, USA).

## 3. Results

### 3.1 Plant identification and protein concentration

The identification of the studied plants was performed by a botanist from the Jardim Botânico do Rio de Janeiro. The plant purchased on the market (Mercadão de Madureira) was identified as *Eugenia astringens* Cambess., of family Myrtaceae, and the plants collected in Marechal Deodoro, Massambaba and Praia Grande as *Chrysobalanus icaco* L., plant from family Chrysobalanaceae.

Chrysobalanaceae can be morphologically diffeenciate from the Brazilian Myrtaceae species, by some characteristics, such as phylotaxia, which is alternating (and opposite in Myrtaceae). Nevertheless, the similar form of the leaves of *C. icaco* and *E. astringens* can cause confusion (Fig. 2), the *E. astringens* leaf has a fold around it facing the abaxial part (Fig. 2c) not found in *C. icaco*.

**Fig. 2.**
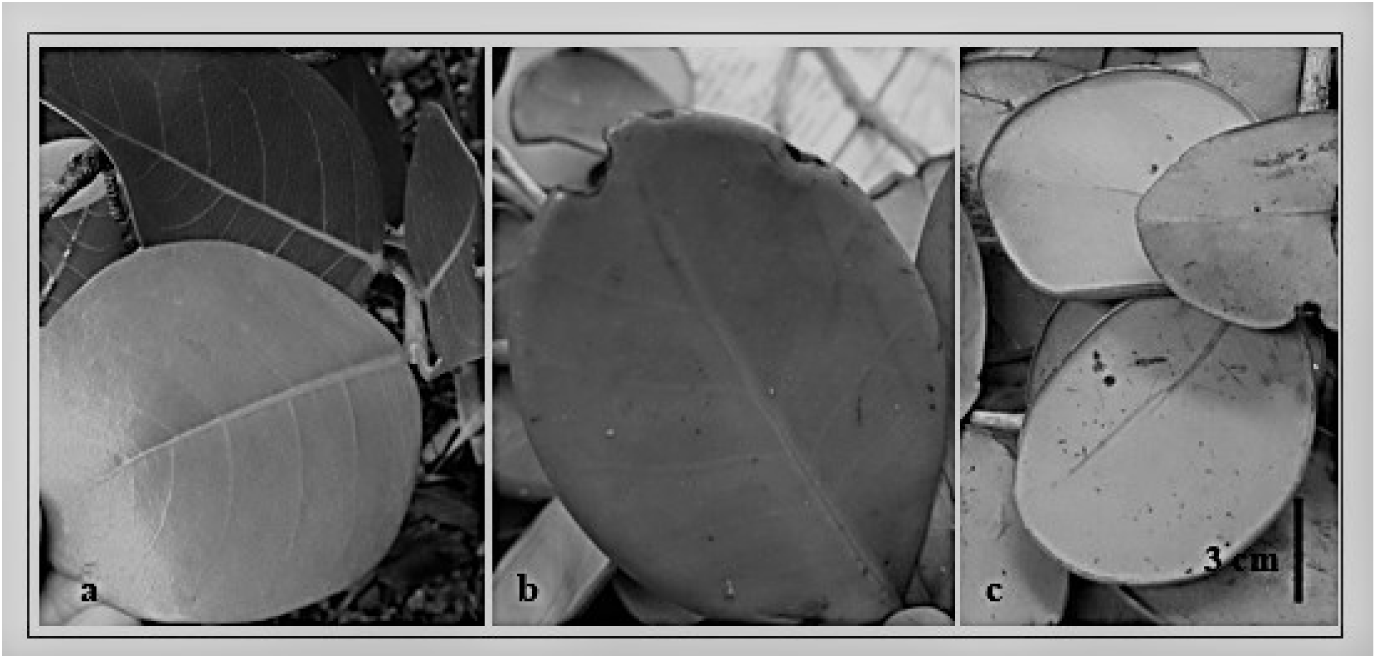
Comparison between the branches of *Chrysobalanus icaco* L. (Chrysobalanaceae) (a) and *Eugenia astringens* Cambess (Myrtaceae) (b). Abaxial part of *E. astringens* leaf (c). Source: Photos by the author.

This misconception has been previously reported (Bochner *et al*., 2012; Silva and Peixoto, 2009), claiming that the trade of medicinal plants is not a safe source of sale, as it may be difficult for both the trader and the consumer to correctly identify a desirable plant. And yet there is the problem that different plants known by the same popular name are commercialized without proof of their pharmacological properties and toxicological safety (Bochner *et al*., 2012), besides the adulteration possibilities. Unfortunately, in Brazil, the supervision of trade of medicinal plants by regulatory agencies is still incipient.

Total protein concentrations ranged from 30.18 to 54.95 μg μL^-1^ in *Eugenia astringens*, from 28.01 to 43.88 μg μL^-1^ in *Chrysobalanus icaco*.

### 3.2 Cytotoxicity evaluation of protein extracts

Fibroblasts (3T3 cell line) were submitted to the cell viability assay, exposed to different concentrations of protein extract and to the cell migration assay, exposed to a determined concentration of this extract.

#### 3.2.1 Cell viability assay

To evaluate the effects of extracts of *E. astringens* (MAD), *C. icaco* (RMA), *C. icaco* (AL) and *C. icaco* (PG) on fibroblast viability, the MTT assay was performed.

The results for the cell viability assay are shown in Fig. 3, in which it can be observed and compared the reduction of fibroblasts viability among the species and protein extract concentration.

**Fig. 3.**
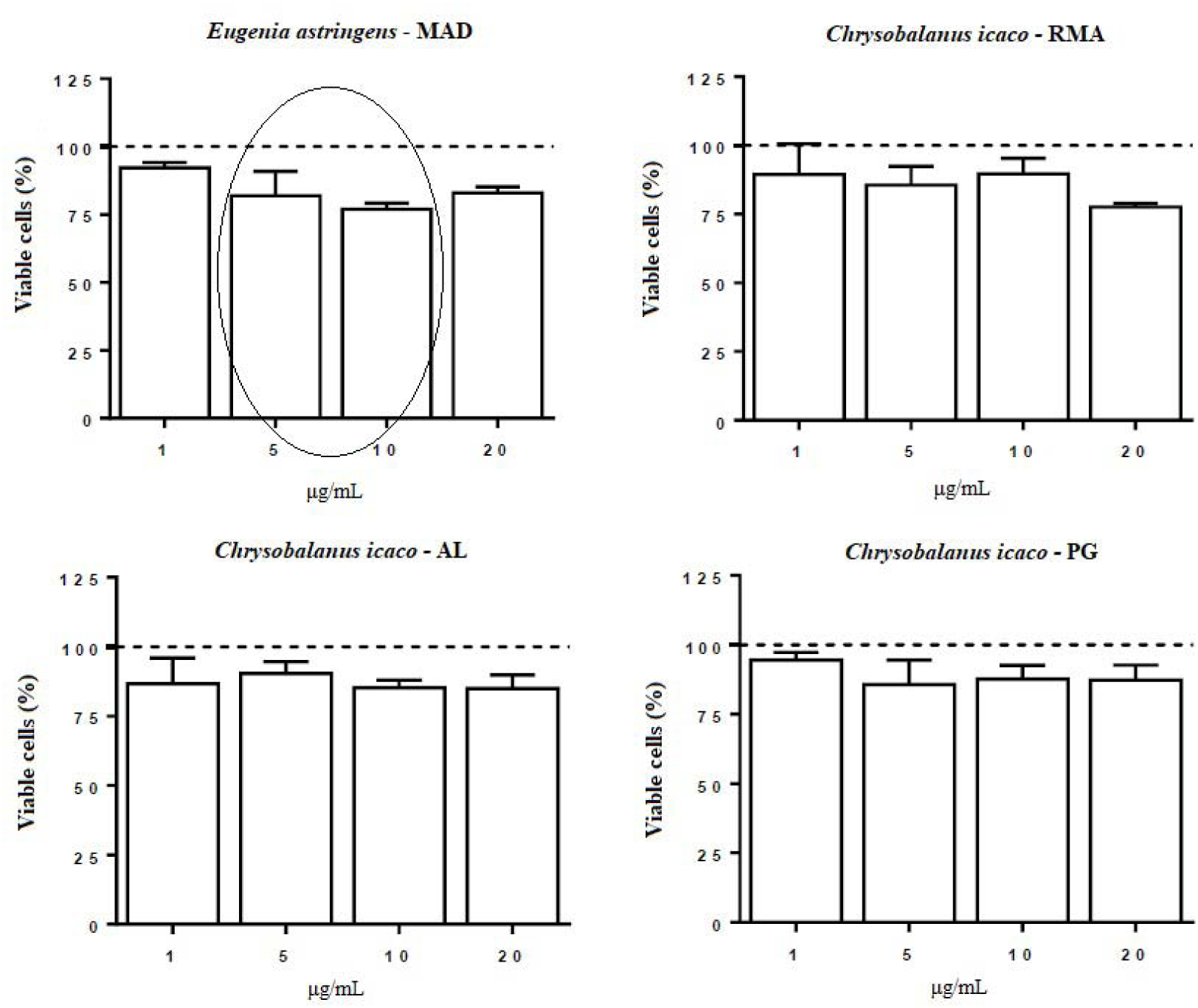
Effect of protein extracts of *Eugenia astringens* and *Chrysobalanus icaco* on fibroblasts viability. MAD – Mercadão de Madureira, RMA - Restinga de Massambaba, AL - Marechal Deodoro, PG - Praia Grande (PG). The dashed line represents the control group (100 %). The circle indicates high reduction on fibroblast viability for *E. astrigens* treatment. Results are mean ± S.E.M. n = 4.

Treatment with *E. astringens* (MAD) at all concentrations tested, reduced cell viability, decreasing by 8.8% (1 μg/mL), 19.2% (5 μg/mL), 23% (10 μg/mL) and 17% (20 μg/mL) the percentage of viable cells. Exposure with *C. icaco* (RMA) at concentrations of 1, 5 and 10 μg/mL did not alter significantly the fibroblasts viability. On the other hand, the increase in concentration resulted in a decrease in the percentage of viable cells, leading to a reduction of 22.4% (P <0.001) in cell viability when the highest concentration (20 μg/mL) was used. Treatment with *C. icaco* (AL), in turn, induced a decrease in cell viability (8-16 %) at all concentrations tested, when compared to the control group. In addition, treatment with *C. icaco* (PG) at 1 μg/mL did not alter the viability of fibroblasts, whereas treatment with the other concentrations induced a decrease in cell viability (12-14 %).

#### 3.2.2 Scratch wound healing assay

To evaluate the effects of extracts of *E. astringens* (MAD), *C. icaco* (RMA), *C. icaco* (AL) and *C. icaco* (PG) on fibroblast migration, the cell migration assay (Scratch Wound Healing Assay) was performed.

As shown in Figure 4, treatment with *C. icaco* (RMA), *C. icaco* (AL) and *C. icaco* (PG) was not able to alter the migration rate of fibroblasts. On the other hand, it can be noted that the treatment with *E. astringens* led to a significant reduction in the migration of these cells by 26.04% (p <0.05), comparing to the control.

**Fig. 4.**
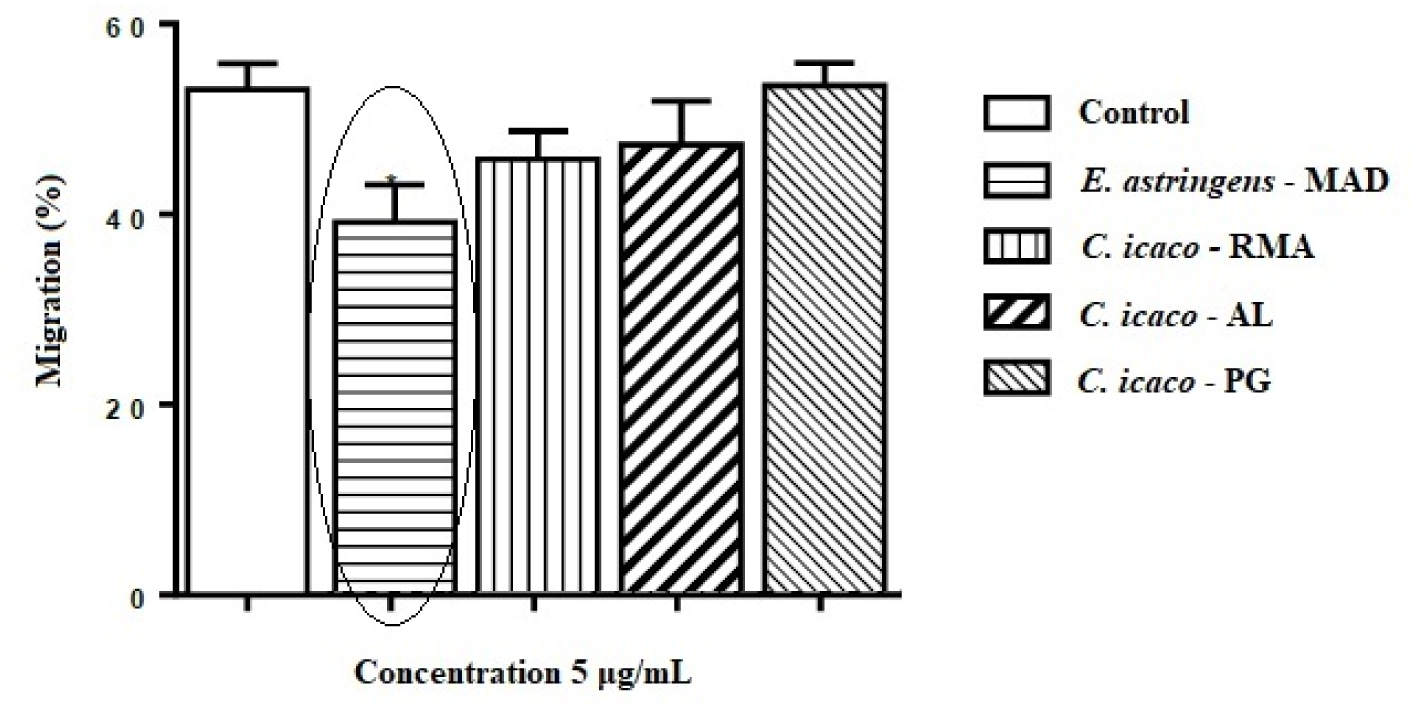
Effect of the extracts of *Eugenia astringens* and *Chrysobalanus icaco* on the migration of fibroblasts at times 0 and 24 hours. MAD – Mercadão de Madureira, RMA - Restinga de Massambaba, AL - Marechal Deodoro, PG - Praia Grande (PG). Circle indicates significant reduction on fibroblast migration for *E. astrigens* treatment. Results are mean ± S.E.M. One-way Anova, followed by Newman-Keuls post-test, *p <0.05. n = 3.

The migration of fibroblast is illustrated in Fig. 5, in which can be observed a lower migration of these cells when treated with *E. astringens* protein extract than the *C. icaco* treatments, slowing wound closure.

**Fig. 5.**
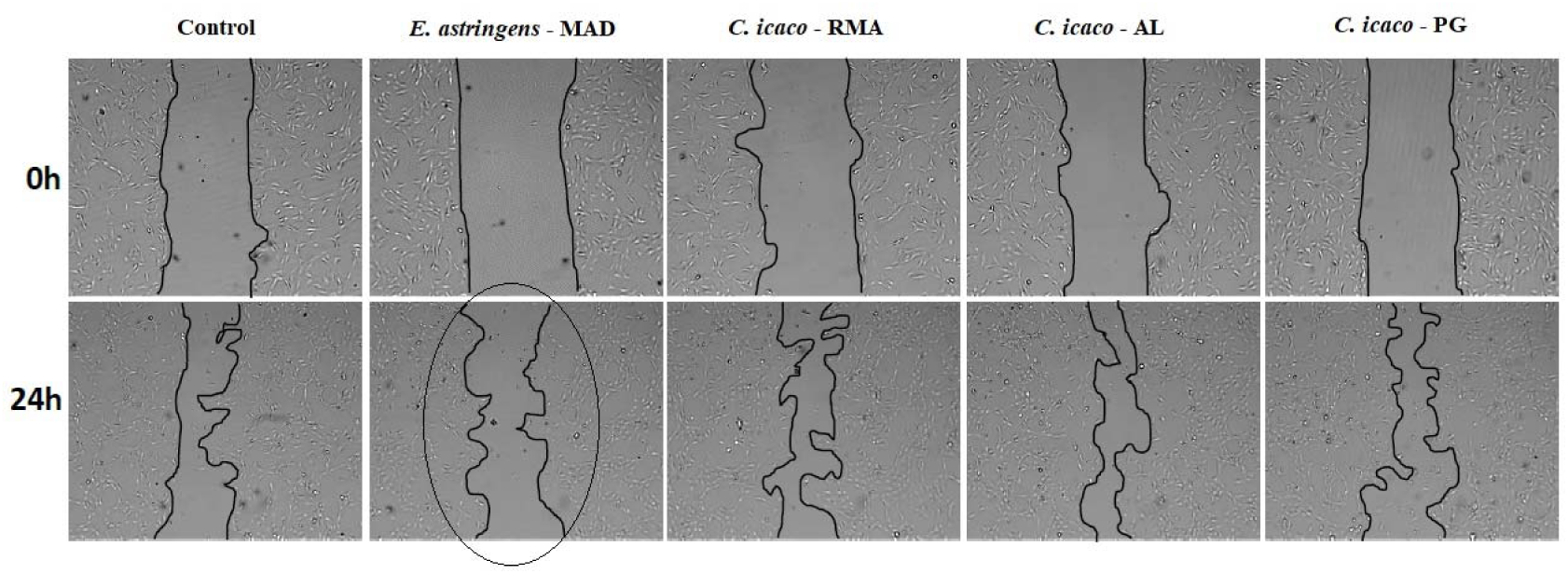
Effect of the extracts of *Eugenia astringens* and *Chrysobalanus icaco* on the migration of fibroblasts (3T3) at 0 h and 24 hours. MAD – Mercadão de Madureira, RMA – Restinga de Massambaba, AL – Marechal Deodoro, PG - Praia Grande. Circle indicates significant reduction on fibroblast migration for *E. astrigens* treatment. n=3.

## 4. Discussion

There is a misunderstanding regarding the sale of abajerú in Mercadão de Madureira, where *Eugenia astringens*, of the same popular name, is sold in place of *Chrysobalanus icaco*. This is of great concern to Public Health, because *C. icaco* is popularly used as medicinal plant for treating diabetes, due to its hypoglycemic potential. Meanwhile the population consumes tea from the leaves of *E. astringens*, coming from these markets instead of *C. icaco*. Medicinal plants are widely used due to their easy accessibility, but they usually not have their efficacy and safety well established (Bochner *et al*., 2012). This fact can become a risk to those who use them since they can cause more deleterious effects than bring health benefits. It is of prime importance to inspect qualified individuals, traders, distributors and producers for regularization of the sale of medicinal plants.

Silva and Peixoto (2009) raised three hypotheses regarding the introduction of *Eugenia astringens*, replacing *Chrysobalanus icaco* in popular marketing. First, it would be a strategy of the merchants to circumvent the competent oversight, by having the same popular name, but neither could it distinguish. A second hypothesis would be related to the difficulty in the recognition of the species by the collectors and sellers, as well as the consumers, due to the similar morphology. The last hypothesis would be the attribution of hypoglycemic activity to *E. astringens* by herbivores, since other species of Myrtaceae, such as pitanga (*Eugenia uniflora* L.), jambo (*Eugenia jambos* L.) and *Eucalyptus*, are used by the population for this purpose and have antioxidant, antifungal and antibacterial properties (Queiroz *et al*., 2015). Also, the natural environments to which *C. icaco* occurs are restinga-type vegetation sites, which are usually areas of environmental protection, which makes it difficult to collect specimens of this species. Therefore, this could also be a hypothesis regarding the introduction of *E. astringens*, replacing *C. icaco* in popular marketing. This species is not hypoglycemic like *C. icaco*, which can lead to intoxication in those people who buy erroneously, thinking that they are acquiring the correct abajerú plant.

Since medicinal plants may also have unknown toxicological properties, the evaluation of toxicity, through in vitro tests, is required. Cytotoxicity of the extracts of medicinal plants, including those that are hypoglycemic, can affect cellular processes like healing that is crucial for diabetic patients. Hyperglycemia alters leukocyte function, increasing the risk of bleeding and impairing inflammatory and healing processes (Negri, 2005; Aquino *et al*., 2019). This difficulty in healing occurs due to cardiovascular complications, which cause blockage or decrease of blood circulation, and due to excess glucose, which can impair the functioning of the immune system. That is, diseased vessels decrease blood flow, especially to the legs and feet, harming the healing process and high glycemic levels incapacitate the body’s defense cells (Hu et al., 2002).

Zandi *et al*. (2016) verified the viability of fibroblasts (ovine line) in extracts of different plants (*Aloe vera*, hena, camomile, licorice, myrtle, mint, cinnamon, ginger and cedar) and that at the minimum concentration (6.25 μg/mL), the viability of dermal fibroblasts by MTT assay increased significantly in cedar (p <0.05). Combination of *Aloe vera*, mint extract and licorice significantly increased the viability of dermal fibroblasts (p <0.05). *Aloe vera*, which is also known for its hypoglycemic activity, has the ability to stimulate proliferation of L929 fibroblasts (Manoj *et al*., 2009). Calloni *et al*. (2016) tested the phenolic extract of *Plinia trunciflora* from the same family as *E. astringens* on human lung fibroblast cells in the presence and absence of amiodarone, a drug used to treat arrhythmia, but which causes toxicity in the lungs. The extract rich in polyphenols was able to prevent the decrease of cellular viability (MTT test) and the ATP biosynthesis.

There are no studies testing the viability of fibroblasts exposed to protein extracts of *Chrysobalanus icaco*, but ethanolic extracts of these species prove to be important in cellular processes. Silva *et al*. (2017) evaluated the antifungal activity of the *C. icaco* ethanolic extract, noting the inhibition of growth of *Candida albicans* and *C. parapsilosis*, strains exposed to this extract.

Pitz *et al*. (2016) evaluated the in vitro activity of ethanolic extract of *Plinia peruviana* bark, the same family as the *E. astringens*, in healing processes and antioxidant activity in urinary fibroblasts (L929 cell line). The cell migration assay (Scratch Wound Healing Assay) indicated that none of the tested shell concentrations (0.5, 5, 25, 50 and 100 μg/ml) was able to increase the migration rate after 12 hours of incubation. These results demonstrate a positive effect of the peel on the wound healing process in the L929 fibroblast cell line, probably due to the antioxidant activity exhibited by phytochemicals in the extract. Manoj *et al*. (2009) verified the effect of germplasm of *Aloe vera*, which is also hypoglycemic in L929 fibroblasts, through the cell migration assay, confirming the increase in fibroblast migration, which is important for regeneration and skin repair in case of injury.

There are no studies testing the viability and migration of fibroblasts exposed to protein extracts of *C. icaco* and *E. astringes*, however ethanolic extracts are used in studies to test toxicity of *Eugenia* species. The in vitro antioxidant activity of the ethanolic extract of *Eugenia uniflora* was determined by the inhibition of spontaneous autoxidation in brain homogenate, with the LD_50_ of 5.93 g/kg in mice (Auricchio *et al*., 2007). In the phytotoxicity test of the *Eugenia catharinae* extract, it was observed that ethyl acetate and hexane fractions inhibited seed germination, while the hexane fraction showed higher inhibition of lettuce seedlings. *E. catharinae* demonstrated a considerable toxic activity, encouraging the search for the compounds responsible for this activity (Colla and Brighente, 2011).

## Conclusion

The assays to evaluate the toxicity of the protein extracts of the plants studied served to make aware of the sale and use of the *Eugenia astringens* plant, sold in place of *Chrysobalanus icaco*, since it reduced cell viability at all concentrations of the extract and decreased the fibroblast migration rate. These results showed that *E. astringens* can cause cytotoxic effects if consumed in larger doses.

The present work demonstrated the importance of research in the area of Public Health and the dissemination and communication to society of the results of scientific works since, due to the confounding of the use of medicinal plants, diabetic patients may opt for natural products in therapeutic use for the treatment of diabetes, in the wrong way.

## Acknowledgements

The authors are thankful to CNPq, to CAPES and to Dra. Viviane Kruel from the Research Institute of Botanical Garden of Rio de Janeiro for the identification of the specimens collected. The cytotoxicity assays were performed at the Laboratory of Cell Biology of the Federal University of Alagoas under the supervision of Dr. Emiliano Barreto. This study was carried out with financial support from the Coordination and Improvement of Higher Level or Education Personnel – CAPES (PhD’s grant).

## Abbreviations

MAD: Mercadão de Madureira;
PG: Praia Grande;
RMA: Restinga de Massambaba;
AL: Marechal Deodoro;
MTT: 3- (4,5-dimethylthiazol-2-yl)-2,5-diphenyltetrazolium bromide.

